# Improved Ultrasound Localization Microscopy based on Microbubble Uncoupling via Transmit Excitation (MUTE)

**DOI:** 10.1101/2021.10.05.463265

**Authors:** Jihun Kim, Mathew R. Lowerison, Nathiya Chandra Sekaran, Zhengchang Kou, Zhijie Dong, Michael L. Oelze, Daniel A. Llano, Pengfei Song

## Abstract

Ultrasound localization microscopy (ULM) demonstrates great potential for visualization of tissue microvasculature at depth with high spatial resolution. The success of ULM heavily depends on the robust localization of isolated microbubbles (MBs), which can be challenging in vivo especially within larger vessels where MBs can overlap and cluster close together. While MB dilution alleviates the issue of MB overlap to a certain extent, it drastically increases the data acquisition time needed for MBs to populate the microvasculature, which is already on the order of several minutes using recommended MB concentrations. Inspired by optical super-resolution imaging based on stimulated emission depletion (STED), here we propose a novel ULM imaging sequence based on microbubble uncoupling via transmit excitation (MUTE). MUTE “silences” MB signals by creating acoustic nulls to facilitate MB separation, which leads to robust localization of MBs especially under high concentrations. The efficiency of localization accomplished via the proposed technique was first evaluated in simulation studies with conventional ULM as a benchmark. Then an in vivo study based on the chorioallantoic membrane (CAM) of chicken embryos showed that MUTE could reduce the data acquisition time by half thanks to the enhanced MB separation and localization. Finally, the performance of MUTE was validated in an in vivo mouse brain study. These results demonstrate the high MB localization efficacy of MUTE-ULM, which contributes to a reduced data acquisition time and improved temporal resolution for ULM.

## I. INTRODUCTION

UPER resolution ultrasound localization microscopy (ULM) is an acoustic analog to optical super-resolution microscopy such as photoactivated localization microscopy (PALM) [1] or stochastic optical reconstruction microscopy (STORM) [2]. It was recently introduced for microvasculature imaging beyond the inherent acoustic spatial resolution limit while conserving the imaging depth of conventional ultrasound. The primary idea of ULM is to localize microbubbles (MBs) flowing in the vascular networks to achieve super-resolution, and then track the localized MBs over time to measure blood flow velocity [3]. ULM improves conventional ultrasound spatial resolution by approximately 10-fold and showed promising results in various tissues including the brain [4], kidney [5], liver [6], and tumor [7]. However, practical implementation of ULM is currently limited by its Achilles’ heel – slow temporal resolution, which is largely attributed to the long data acquisition time that is required to capture adequate, spatially separated MB signals for localization and tracking. Although increasing MB concentration alleviates slow MB perfusion and shortens data acquisition time, it causes more MB signals to overlap and become unlocalizable. As such, developing a method that is highly efficient at localizing MBs under high concentration is essential to accelerate ULM.

To this end, several techniques have been demonstrated to improve the super-resolution imaging speed, including super-resolution optical fluctuation imaging (SOFI)-based contrast-enhanced ultrasound (CEUS) imaging [8], and sparsity-based ultrasound super-resolution hemodynamic imaging (SUSHI) [9]. SOFI-based CEUS imaging utilizes the high order statistics of the fluctuating MBs signal. In contrast, SUSHI exploits the sparsity model by assuming the underlying vasculature consists of point targets on the higher pixel resolution. These techniques showed improvement of temporal resolution while providing a spatial resolution that is similar to ULM. However, because of the absence of MB tracking, these methods fall short of providing blood flow velocity measurements, which can be an important biomarker for various applications [4], [10], [11].

Recently, our group introduced a MB separation method that separates the high concentration MBs in the Fourier domain to facilitate localization of MBs under high concentration [10]. Thanks to the enhanced localization efficacy, MB separation leads to a reduced data acquisition time for ULM. However, by creating subsets of data with different MB flow characteristics, MB separation becomes computationally expensive. In addition, the performance of MB separation may be undermined by complex flow dynamics with high MB concentrations. Compressed sensing-based localization techniques also showed the capability of reducing data acquisition time for ULM under MB high concentration, but it has a similar issue of high computational cost as the MB separation method and it also does not provide blood flow velocity measurement [12].

Inspired by the optical super-resolution technique STED (stimulated emission depletion), in this paper, we propose to implement a similar principle of contrast signal suppression in ultrasound imaging to promote MB separation and enhance ULM imaging performance. STED employs two transmissions including a Gaussian-shaped beam followed by a doughnut-shaped beam that depletes the fluorescence excited by the Gaussian beam. The partial depletion preserves a sharp focal spot in the donut center, where a small number of fluorophores are allowed to emit signals that are distinct from surrounding fluorophores that have been depleted [13]. The distinction between the fluorophores inside and outside the donut center creates separation in fluorescence status in time, which manifests as super-resolution when imaging.

While the principle of STED is straightforward, translating it to ultrasound imaging is not because in ultrasound one needs to overcome the challenge of depleting MB signals, which is not as simple as in optics. For example, to deplete MB signal, one needs to apply high mechanical index (MI) acoustic pulses to disrupt MBs, which is not an efficient process and may cause tissue and probe heating. Also, unlike in optical STED where a fluorophore can be instantly and completely switched to the “off” or ground status by the stimulation photon, MB signal emission is less binary (e.g., either “on” or “off”) and even disrupted MBs can emit strong acoustic signals [14]. As such, an alternative approach is necessary for ultrasound imaging to implement the principles of STED.

In this paper, we introduce a method named MUTE (microbubble uncoupling via transmit excitation) that uses the principles of acoustic null generation and subtraction imaging to realize STED in ultrasound. MUTE accomplishes MB depletion in two steps: 1) obtain two images by first transmitting a regular plane wave (PW) without acoustic nulls and then another PW with acoustic nulls; 2) calculate the difference between the two images and then MBs in the non-null region will be effectively depleted and MBs in the null region will be enhanced and separated from the ones in the non-null region. Although similar ideas of using acoustic nulls to enhance ultrasound imaging resolution have been reported before, they were implemented on receive apodization which does not actively modulate the acoustic response of MBs by transmit excitation [15], [16]. In addition, these methods suffer from a low signal-to-noise ratio (SNR) due to the significantly reduced signal intensity induced by the processes of Heaviside apodization and subtraction [17]. With MUTE, one can create a similar on/off status of MBs as in STED, which provides the opportunity to spatially separate MBs that are otherwise inseparable under traditional insonification. By promoting MB separation, MUTE facilitates robust MB localization for ULM, especially under high MB concentrations. The improved MB localization efficacy leads to an overall reduced data acquisition time for ULM.

The rest of the paper is structured as follows: MUTE imaging sequence, post-processing steps, and preparation of models for evaluation including simulation, chorioallantoic membrane (CAM) of a chicken embryo, and a mouse brain are presented in Section II. In Section III, we will demonstrate the improvement of MUTE-based ULM imaging with conventional ULM as reference. Finally, our results are discussed, and conclusions drawn in Section IV.

## II. Materials and methods

### A. Principles of Microbubble Uncoupling Via Transmit Excitation (MUTE)

Figure 1(a) shows the conceptual schematic of MB uncoupling in tissue vasculature based on MUTE. Flowing MBs in vessels are activated or silenced depending on their spatial location with respect to the acoustic null. By calculating the difference between two consecutive images (i.e., an image with the acoustic null and the other without), MBs in the non-null region will be suppressed, and MBs in the null region will be enhanced and subsequently uncoupled from surrounding MBs. In general, images with acoustic nulls can be generated by apodizing the transmit aperture with functions such as the Heaviside [15], for example:

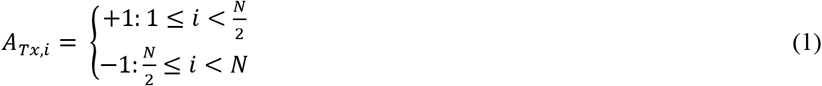

where *A_Tx,i_* is the transmit apodization weight for the *i th* element, and *N* is the total number of active elements in the transmit aperture. Figure 1(b) shows the beam pattern of PW (left) and MUTE (right) acquired by adjusting the transmit apodization weight. The left and middle panel of Fig. 1(c) shows B-mode images of 3 MBs spaced with 2λ distance (λ is wavelength), based on PW transmission and MUTE transmission, from which uncoupled MB signals can be obtained by calculating the difference between the two images. To minimize image distortion by subtraction, an intensity weighted subtraction algorithm was adopted for the MUTE sequence [18]:

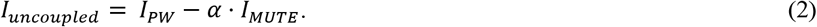

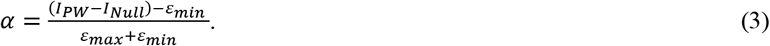

where *I_PW_* and *I_MUTE_* are the intensity images (i.e., B-mode) obtained by PW and MUTE, respectively. *ε_max_* and *ε_min_* are the upper and lower bounds of the intensity values of the uncoupled MB image, in which we set −1 and +1, as the minimum and maximum intensity values. The bottom-row in Fig. 1(c) shows the normalized intensity profiles in the lateral direction corresponding to each B-mode image. One can clearly observe that the center MB in the acoustic null that was not distinguishable from the surrounding MBs now becomes distinct and localizable for ULM imaging.

**Fig. 1.**
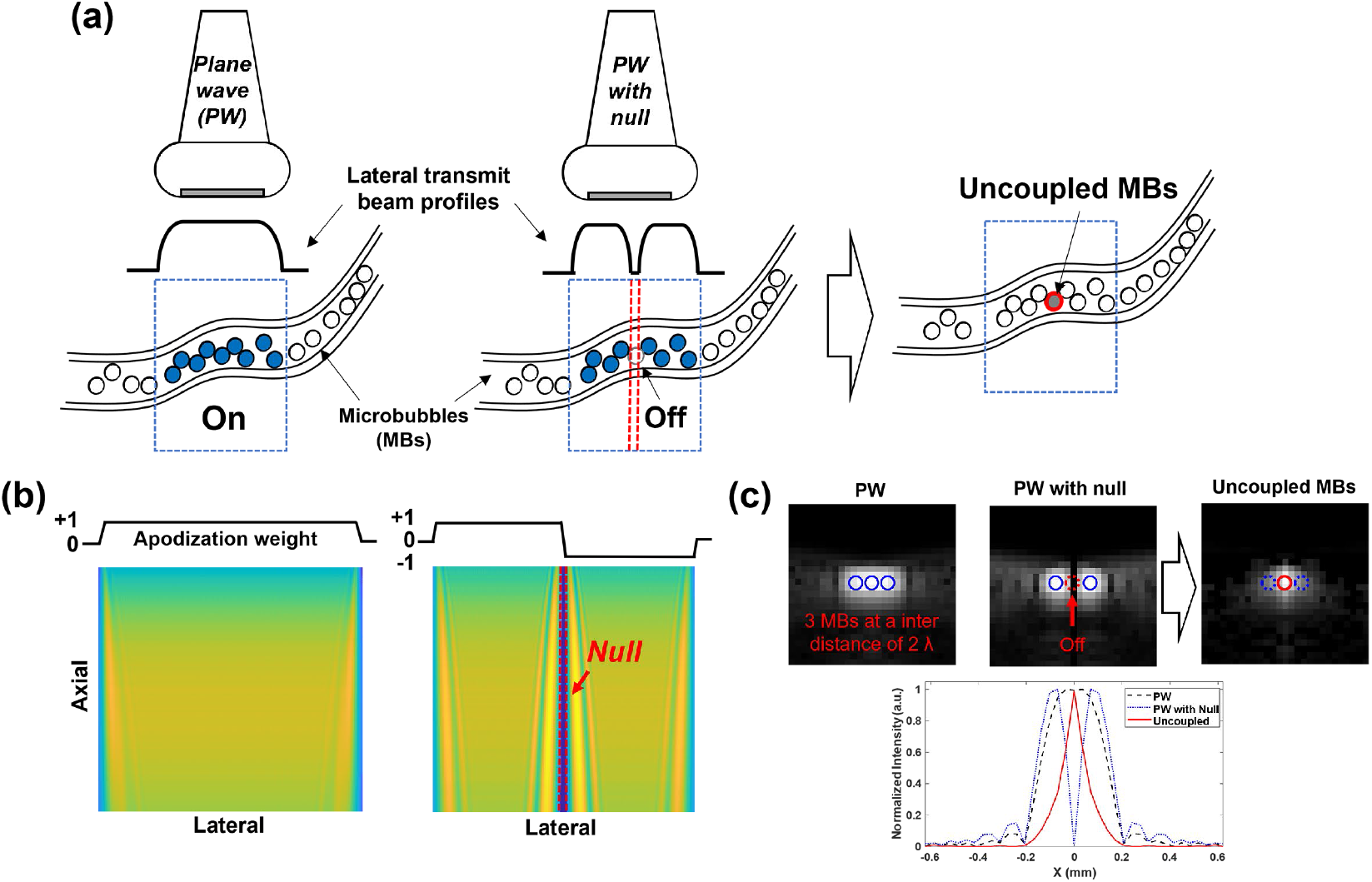
Overall concept of MUTE imaging: (a) Schematic for MBs uncoupling, (b) Beam patterns of a PW (left) and PW with null (right) by adjusting the transmit apodization weight, and (c) B-mode images of 3-MBs spaced with 2λ distance (λ is wavelength), based on PW (left) and PW with null (middle). Uncoupled MBs obtained by calculating the difference between the PW image and the PW with null image. The rigid and dashed circles in (c) represent the “On” and “Off” status of MBs, respectively. The bottom-row in (c) shows the normalized intensity profiles in the lateral direction corresponding to each B-mode image in the top-row.

### B. MUTE Imaging Sequence and Data Acquisition

In practice, because a single acoustic null only uncouples MBs in a small-limited region, it becomes challenging to implement MUTE with an adequate frame rate for ULM imaging (e.g., one needs to translate the null to different lateral locations in different pulse-echo cycles). Also importantly, since a single PW does not provide transmit focusing and adequate imaging SNR, it is necessary to integrate spatial angular compounding into the MUTE sequence for practical use [19]. To address these practical issues, as shown in Fig. 2, we designed a pulse train of Heaviside functions for transmit apodization, which was then steered with various angles to realize spatial compounding by adjusting the transmit delay for each element. In detail, the MUTE beam includes a total number of 31 nulls with equal distances of 5.6λ in the lateral direction. The width of null is 0.7λ. Furthermore, to cover the entire imaging field by nulls, the MUTE beam was translated laterally with a step size of 1.4λ corresponding to the pitch size of the transducer we utilized, resulting in the use of the four MUTE beams at each steering angle. To minimize overlap of non-null regions between different steering angles while still providing ample angular compounding, a step steering angle of 2 degrees was chosen throughout the study. As shown in Fig. 2, upon spatial compounding of each MUTE sequence, the acoustic nulls were preserved while the non-null regions were compounded with enhanced transmit focusing and SNR.

**Fig. 2.**
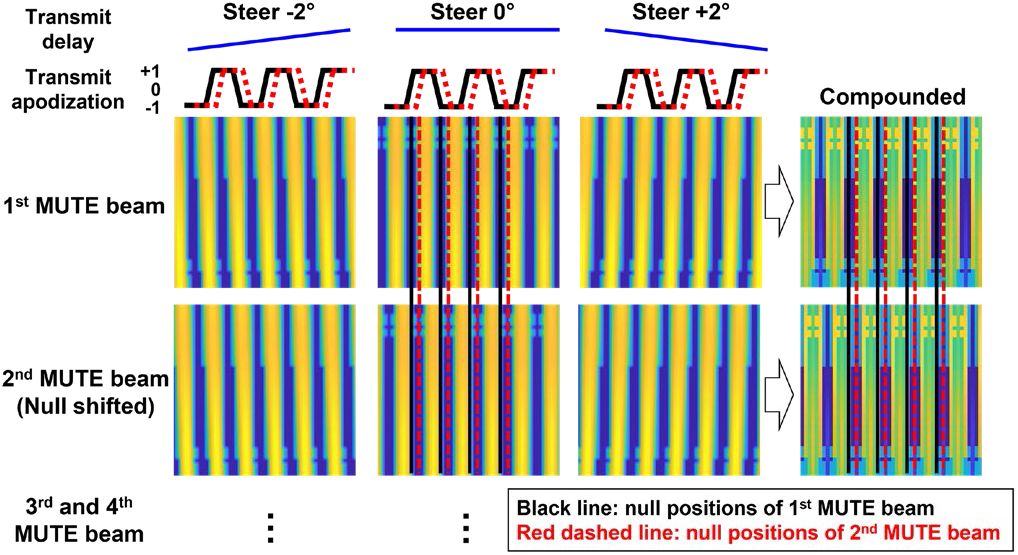
MUTE imaging sequence. Beam patterns of four MUTE sequences with three steering angles of −2°, 0 °, +2°. Each MUTE sequence has different acoustic null positions as indicated by the black and red dashed line depending on the transmit apodization shifting.

For the acquisition of ultrasound in-phase and quadrature (IQ) data, we utilized an ultrasound imaging system (Vantage 256, Verasonics Inc., Kirkland, WA, USA) with a high-frequency linear array transducer (L35-16vx, Verasonics Inc., Kirkland, WA, USA). The receiving demodulation frequency was set as 31.25 MHz. The transmit frequency of 20 MHz with a single sinusoidal wave and transmit voltage of 10 V were utilized. Furthermore, ultrasound imaging was performed using three steering angles with a step steering degree of 2 (−2°, 0°, +2°). PW and four MUTE beams were sequentially generated at each steering angle and coherent compounding was performed separately to each IQ sub-datasets. The post-compounded effective frame rate was 500 Hz. The total number of 5,000 IQ datasets including each 1,000 IQ sub-datasets for PW and four MUTE sequences were stored per one acquisition, which corresponds to the data acquisition time of 2s.

To measure the beam pattern and MI generated by the MUTE sequence, a needle hydrophone (1603, Precision Acoustic Ltd., Dorset, U.K.) with a customized acoustic intensity measurement system was utilized.

### C. Post-processing Steps for the Reconstruction of MUTE-ULM image

For the reconstruction of the MUTE-ULM image, the post-processing steps were shown in Fig. 3. A spatiotemporal singular value decomposition (SVD) filter was used to extract the MB signal from surrounding tissue [20], [21]. A noise equalization process was applied throughout the imaging field of view to facilitate robust thresholding based on MB signal intensity [22]. MB separation was used in this study with three filter bands of 15 to 50, 50 to 100, and 100 to 200 Hz [23]. After MB separation, a spatiotemporal non-local means (NLM) filter was used to further reduce noise that may result in false MB localizations [24]. We then performed the MUTE image subtraction based on Eq. (2) followed by MB localization and tracking. For MB localization, MB images were spatially interpolated to 0.1λ pixel size (4.9 μm). Two point-spread functions (PSF) were then constructed for the PW and MUTE sequences, respectively, using a 2D multivariate Gaussian model:

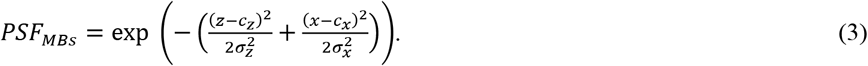

where (*c_z_*, *c_x_*) represents the center location of MB. *σ_z_* and *σ_x_* were calculated in the axial and lateral directions, respectively, as follows:

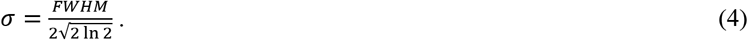

where the full width at half maximum (FWHM) of MBs was measured by averaging 10 random MB signals obtained at different imaging depths and frames. In this study, the average FWHM of MBs via PW in the lateral and axial directions were measured to be 138 μm and 86 μm, respectively; and120 μm and 86 μm for MUTE.

**Fig. 3.**
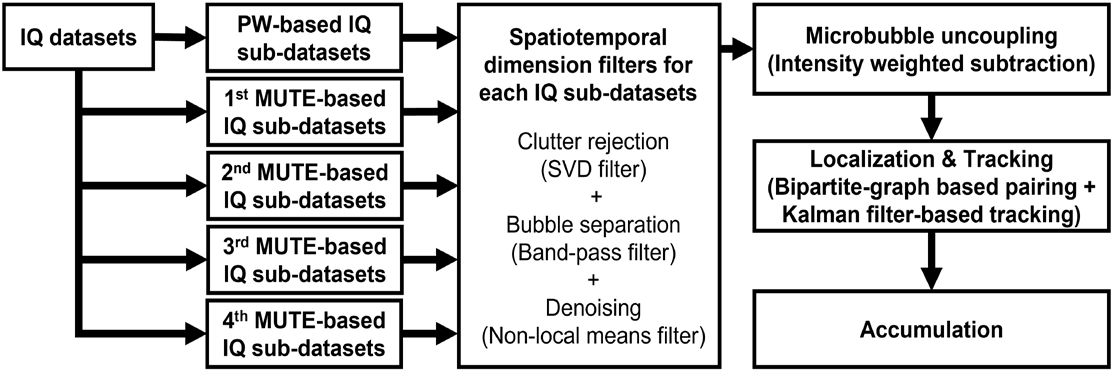
Post-processing steps for reconstruction of MUTE-ULM image including clutter rejection, MB separation, denoising, MB uncoupling, localization, and tracking. These processes were repeated for each imaging frame, and then the final ULM image was reconstructed by an accumulation of MB tracks.

After constructing the appropriate PSFs, a similar process of MB localization and tracking as in our recent papers [24], [25] were performed.

### D. Simulation Study

To evaluate the efficiency and error of localization depending on the concentration of MBs, we produced digital phantoms with randomly distributed MBs at the concentrations of 5, 25, 50, 100, 125, and 150 MBs/mm^2^ using the Verasonics simulator. Furthermore, five phantoms were generated at each concentration to measure the average performance with the standard deviations. The same MUTE imaging sequence as described in Fig. 2 was used in the simulation study. Besides, the same post-processing steps were applied except for the spatiotemporal dimension filters. The localized MBs were paired within 1λ distance with the true MBs generated in simulation using a bipartite graph-based pairing algorithm [24]. Then, the localization efficiency was measured by counting the number of pairings at each concentration. Simultaneously, the localization error was calculated by measuring the distance between paired MBs.

### E. Chicken Embryo Chorioallantoic Membrane (CAM) study

In this part of the study, we used the CAM microvessel with optical imaging ground truth to test the imaging performance of MUTE. Fertilized chicken eggs (white leghorn) were obtained from the University of Illinois Poultry Research Farm and stored in a 37°C egg incubator (Digital Sportsman Cabinet Incubator 1502, GQF Manufacturing Inc., GA, USA). On the fourth day of embryonic development (EDD-04), the eggshells were peeled using a rotary Dremel tool with a cutting wheel, and embryos were transferred into weigh boats. Shell-less embryos were stored in a humidified incubator (HH09, Darwin Chambers Company, MO, USA) until the sixteenth day of embryonic development (EDD-16) to perform the ultrasound imaging.

FDA-approved lipid shell MBs (Definity^®^, Lantheus Medical Imaging Inc., N. Billerica, MA, USA) were used throughout the *in vivo* imaging experiment. The final solution of MBs had a concentration of 1.2 × 10^10^ [26]. For MB injection, a high-order vein on the CAM was cannulated using a glass capillary needle to allow MBs bolus injection of 70 μL intravenously. Then, the ultrasound transducer was placed at the side of the weigh boat. An optical image of the CAM vessels was acquired using a stereomicroscope (SMZ800N, Nikon Corp., Tokyo, Japan) to provide the reference ground truth for imaging performance evaluations. For reconstruction of the ULM image of CAM, a total of 20 IQ datasets corresponding to 20,000 frames were acquired based on the ultrasound imaging setup we described in the previous sections.

To measure the ULM imaging performance between PW and MUTE sequences, a normalized cross-correlation (NCC) was computed between the ULM image and the optical image and used as a quantitative measure of similarity. To accomplish this, the optical image was first resized according to the dimension of the ULM image and then converted to grayscale. Then, a target patch with the size of 98 μm × 98 μm (20 pixels × 20 pixels) was selected from the optical image as the reference. A smaller patch (24.5 μm × 24.5 μm, or 5 pixels × 5 pixels) was then selected from the ULM image for the NCC calculation This process was repeated for the entire imaging FOV (patch translation step size = 4.9 μm, or 1 pixel) and the average NCC coefficients was used as the quantitative measurement of ULM imaging performance. In addition to similarity, we measured the vessel filling (VF) percentage for each imaging sequence. VF is defined as:

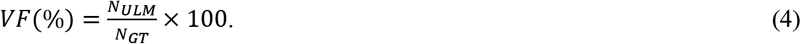

where *N_GT_* denotes the total number of true vessel pixels and *N_ULM_* is the total number of pixels that have been correctly filled by ULM using MB locations. To obtain *N_GT_*, Otsu’s thresholding was applied to the grayscale optical image followed by image dilation and erosion to extract the vascular morphometric information.

### F. In Vivo Mouse Brain Study

All experimental procedures on the mouse presented in this manuscript were approved by the IACUC at the University of Illinois Urbana-Champaign (Protocol number: 19063). Anesthesia was induced using 4% isoflurane mixed with medical oxygen in a gas induction chamber. The mouse was then transferred to a nose cone supplying 2% isoflurane with oxygen for maintenance, and its head was secured to a stereotaxic imaging frame with ear bars. The scalp was removed with surgical scissors, and a cranial window on the left side of the skull was opened with a rotary Dremel tool to expose a hemisphere of the brain. The ultrasound transducer was fixed to a customized stereotaxic imaging stage and positioned above the cranial window to image the brain with a coronal imaging plane. A 30-gauge catheter was then inserted into the tail vein of the mouse for MB injection (50 μL per bolus). Additional 50 μL booster injections were provided throughout the imaging session to maintain the MB concentration in the cerebral vasculature. For reconstruction of the ULM image of the mouse brain, a total of 40 IQ datasets corresponding to 40,000 frames were obtained based on the ultrasound imaging setup we described in the previous sections during two bolus injections.

## III. Results

### A. Measurement of the Beam Pattern and Mechanical Index of MUTE Sequence

We measured the acoustic null width, shift, and distance between the nulls of a MUTE beam using a needle hydrophone. The MUTE beam was generated with a transmit frequency of 20 MHz and transmit voltage of 10 V and at a steering angle of 0°. Fig. 4(a) shows the 2D MI image of the 1st MUTE beam. Every 31 null was detected with an inter distance of 280 μm in a lateral direction corresponding to 5.68λ as we designed the MUTE beam. The maximum MI was measured to be 0.14. Fig. 4(b) represents the lateral MI profiles acquired from the 1st (Black) and 2nd (Blue) MUTE beams at 5 mm in depth indicated by the white dashed line in Fig. 4(a). The width of null was measured at the MI of 0.04 indicated by a gray dashed line in Fig. 4(b). The average width of nulls was measured to be around 50 μm, which corresponds to approximately 1λ. Furthermore, it was confirmed that null locations of the MUTE beam could be laterally moved 70 μm, which corresponds to 1.42λ [Fig. 4(b)].

**Fig. 4.**
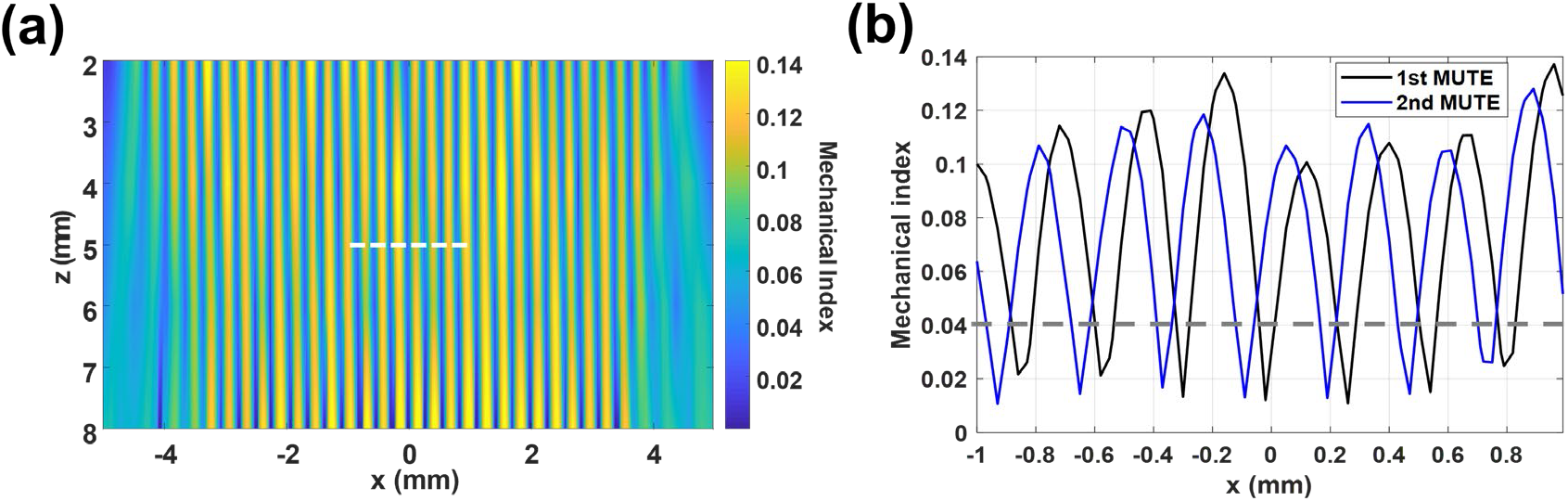
Measurement of the beam pattern and MI of MUTE beam with a transmit frequency of 20 MHz and transmit voltage of 10 V at steering angle of 0° using a needle hydrophone: (a) Representative MUTE beam pattern, and (b) MI profiles of 1^st^ and 2^nd^ MUTE beams acquired at a depth of 5 mm, as indicated by the white dashed line in (a). The gray dashed line in (b) indicates the base line for measuring the width of null. Nulls were detected every 280 μm with a width of 50 μm in the lateral direction. The nulls were shifted laterally with a step size of 70 μm.

### B. Evaluation of MUTE-based Localization Efficacy in Simulation

We evaluated the performance of the PW and MUTE-based localization using different concentrations of simulated MB signals. Figs. 5(a) and 5(b) show the representative localization results based on each dataset obtained by PW and MUTE imaging sequences at the concentration of 50 MB/mm2. The MUTE-based localization had a higher MB detection rate than the PW-based localization, especially for the overlapped MBs indicated by yellow arrows in Figs. 5(a) and 5(b). The average number of MB localizations was 26.4 ± 1.96 MB/mm^2^ for PW, and 41.4 ± 1.74 MB/mm^2^ for MUTE at the concentration of 50 MB/mm2 [Fig. 5(c)]. MUTE-based localization had higher efficacy with a similar localization error across all the MB concentrations tested [Figs. 5(c) and 5(d)]. In particular, as shown in Fig. 5(c), MUTE could localize over 80% of MBs up to 50 MB/mm2 concentration, whereas PW fell short of reaching 80% beyond 25 MB/mm^2^. Moreover, MUTE was largely capable of keeping up with the elevated MB concentration while PW plateaued beyond 50 MB/mm^2^ concentration [Fig. 5(c)]. At the maximum concentration of 150 MB/mm^2^, MUTE was still able to localize over 50% of the microbubbles (81.8 MB/mm^2^). The maximum localization error for both PW- and MUTE-based localization was below 24.64 μm, which corresponds approximately to 0.5λ of the imaging frequency [Fig. 5(d)].

**Fig. 5.**
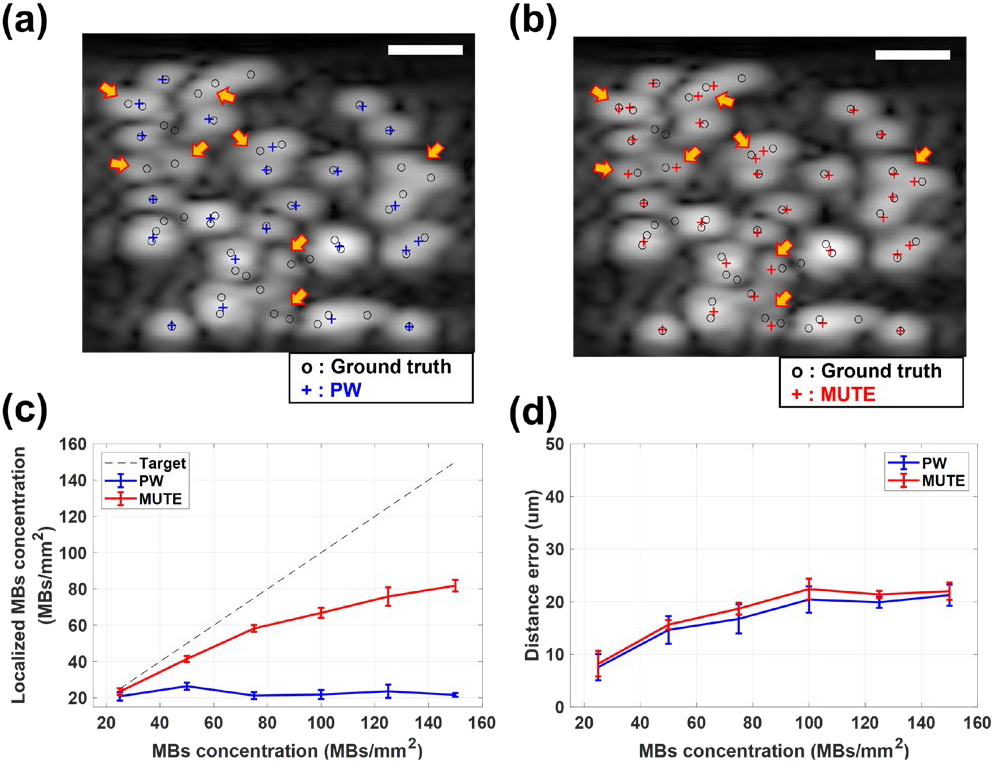
Evaluation of localization efficacy and error using the randomly distributed MBs with different concentrations in simulation: Localization results at the concentration of 50 MBs/mm^2^ overlaid on the B-mode images based on (a) PW and (b) MUTE. Blue and red crosses indicate the localization results of MBs based on PW and MUTE, whereas the black circles indicate the true positions of MBs. (c) and (d) show the localization efficacy and localization error with different concentrations of MBs, respectively. The gray dashed line in (c) indicates the targeted localization efficacy. Scale bars in (a) and (b) represent 200 μm.

### C. MUTE-ULM Imaging of CAM and Quantitative Analysis with Optical Microscopy

The optical image of the CAM in grayscale is shown in Fig. 6(a) and the corresponding binary image generated by Otsu’s thresholding is shown in Fig. 6(b). PW- and MUTE-ULM images are shown in Figs. 6(c) and 6(d), respectively. These images were reconstructed using 20,000 frames of ultrasound data corresponding to a total acquisition time of 40s. Figure 6(e) shows the NCC coefficient between ULM and the optical ground truth versus data acquisition time. It can be observed from Fig. 6(e) that MUTE-ULM generated images with greater similarity to the optical ground truth than did the PW-ULM. The maximum NCC coefficient of PW-ULM was 0.59 at the acquisition time of 40s, while MUTE-ULM achieved the same level of similarity with approximately half the time (22s). Figure 6(f) shows the VF percentage of the two techniques and MUTE-ULM again outperformed PW-ULM across all data acquisition times. MUTE-ULM filled the vessels about twice as fast as did PW-ULM (i.e., 20s vs. 40s) and was able to reach a higher vessel filling percentage than PW-ULM (85.5% vs. 79.9%). Figure 7 shows the local analysis results of the ULM images at the yellow box in Fig. 6. As indicated by the different colored arrows in Figs. 7(a) and 7(b), MUTE-ULM demonstrated faster vessel filling speed and higher overall MB count [Fig. 7(c)] than PW-ULM. MUTE-ULM could detect small vessel signals starting from 2s, while PW-ULM could not detect the small vessels properly at the same acquisition time. As indicated by yellow arrows in Fig 7(c), the 2nd vessel was only detectable with MUTE-ULM and remained missing with PW-ULM until 40s. Moreover, the 4th vessel indicated by the blue arrow was detected by MUTE-ULM at t = 10s [Fig. 7(c)], detected the same vessel at t = 20s. Furthermore, we compared the diameter of the four vessels that were measured at 40s as labeled in Fig. 7(c). Based on MUTE-ULM, the diameter of vessels was measured to be 40.6, 29.7, 52, and 32 μm, respectively, for vessels 1 to 4, while the diameter of vessels was 39.23, 0 (missing), 50, and 30.77 from PW-ULM. The diameters measured from the optical ground truth image were 36.8, 25.6, 42.7, and 28.8 μm. The average error in diameter measurement was 5.1 μm for MUTE-ULM and 9.33 μm for PW-ULM.

**Fig. 6.**
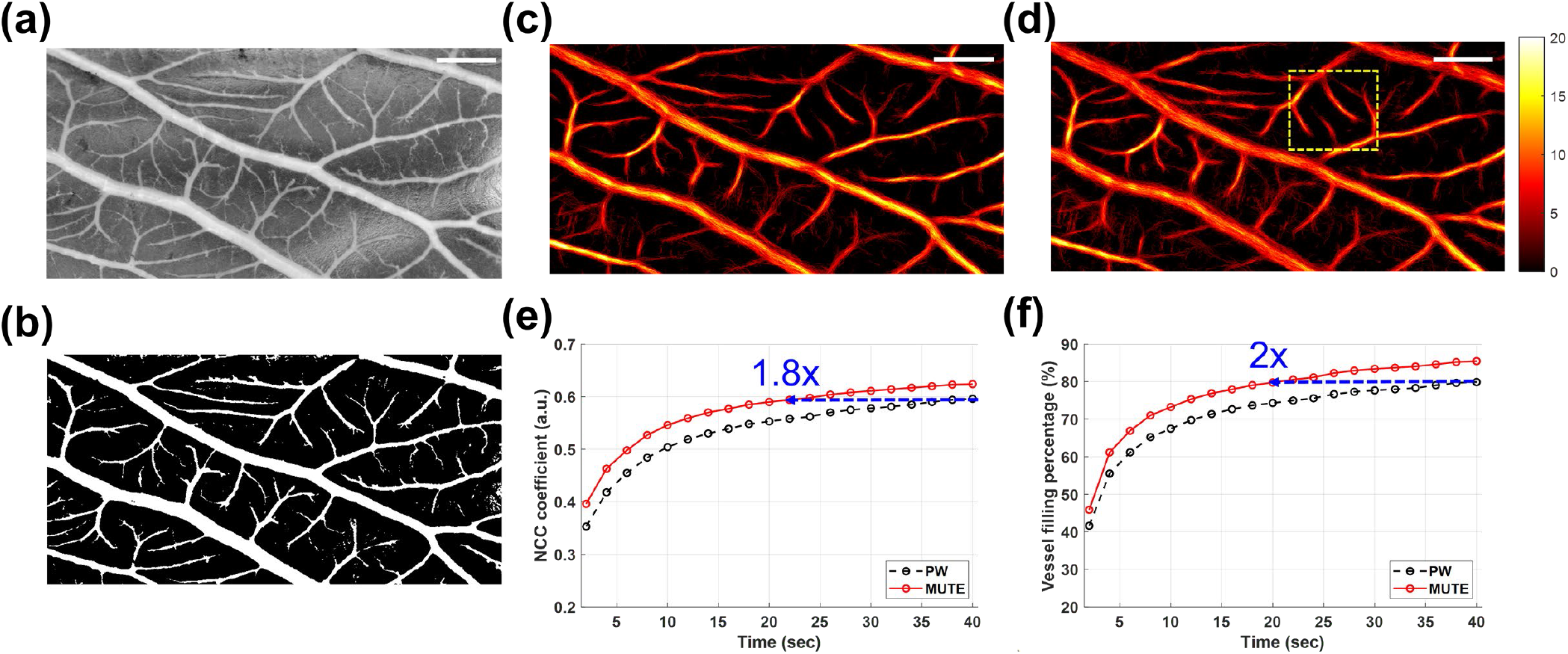
ULM imaging of CAM vessels of chicken embryo validated with optical microscopy: (a) Grayscale optical image, and corresponding (b) Binary optical image, (c) PW- and (d) MUTE-ULM images of CAM vessels. ULM images were represented by the same color scales. (e) Similarity, and (f) VF percentage of each ULM image according to the data acquisition time, validated using the optical binary image of CAM vessels. Scale bars represent 500 μm.

**Fig. 7.**
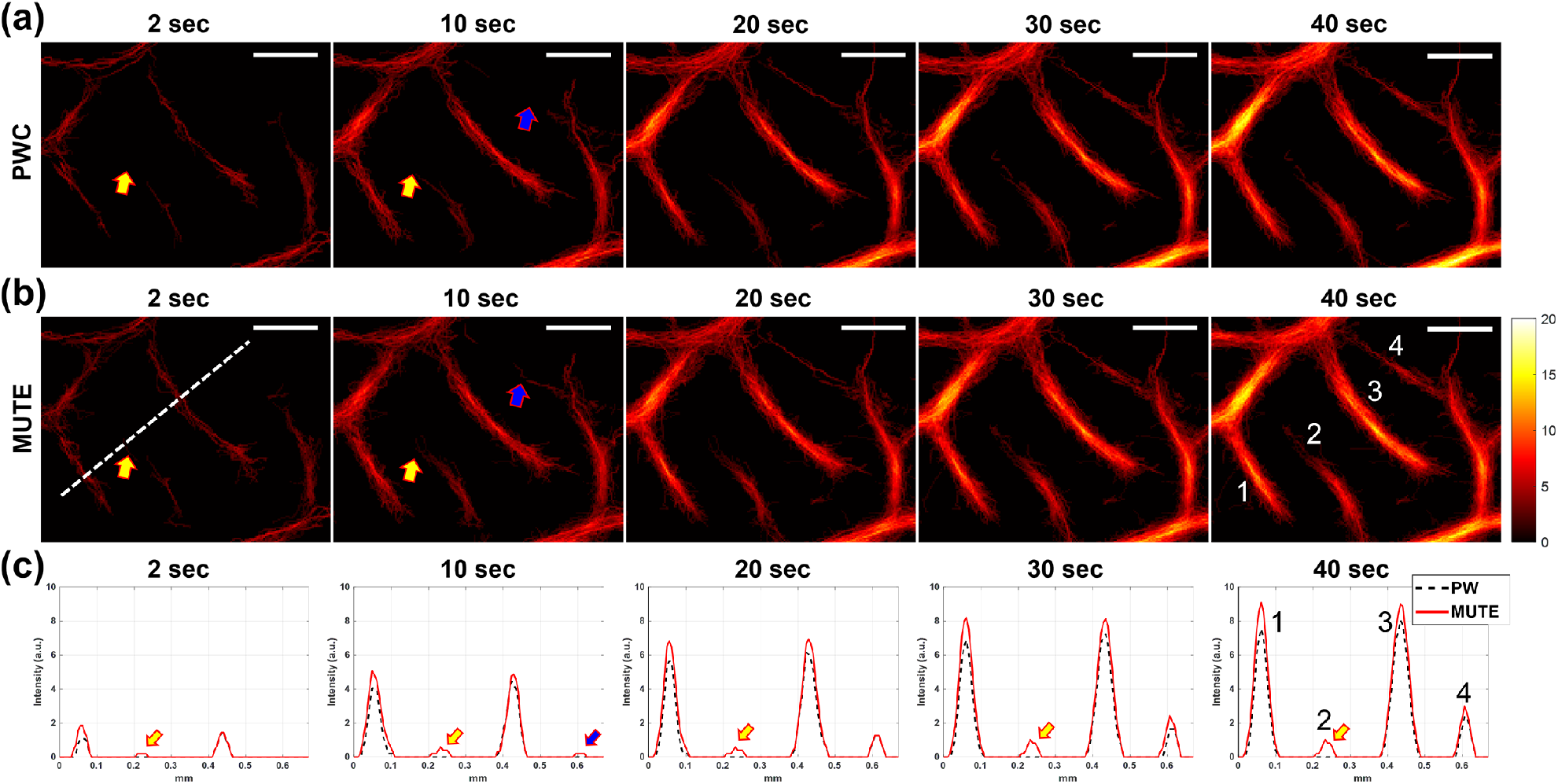
Local analysis of (a) PW- and (b) MUTE-ULM images. The images were extracted from the regions indicated by the yellow dashed box in Fig. 6(d). (c) 1D plots of the vessel signal at different accumulation times from the location indicated by the white dashed line in (b). The different colored arrows indicate different examples of vessels that demonstrate the faster filling speed of MUTE-ULM. Scale bars represent 200 μm.

### D. MUTE-ULM Imaging of the Mouse Brain In Vivo

Figure 8 shows the ULM imaging results using MUTE and conventional PW on a mouse brain. A total of 40,000 frames (80 s) of ultrasound data were acquired during the experiment. We also carried out the local analysis at the regions indicated by the white box in Fig. 8(a). From Fig. 8, both PW- and MUTE-ULM were able to produce robust, super-resolved brain microvasculature maps throughout the full depth of the brain. MUTE-ULM showed higher intensity microvessel density maps that indicate higher MB count, which is most conspicuous in regions with smaller vessels between the large vessels. Similar to the observations from the CAM study, MUTE-ULM showed a faster vessel filling speed than PW-ULM, as indicated by the yellow arrows in Figs. 8(c) and 8(d). The intensity profiles of vessels through the white dashed line indicated in Fig. 8(d) demonstrate that MUTE-ULM detected a higher MB count than PW-ULM, which is again consistent with the results in the CAM study. These results, thus, demonstrate that the proposed technique has the potential to shorten the data acquisition time for in vivo applications.

**Fig. 8.**
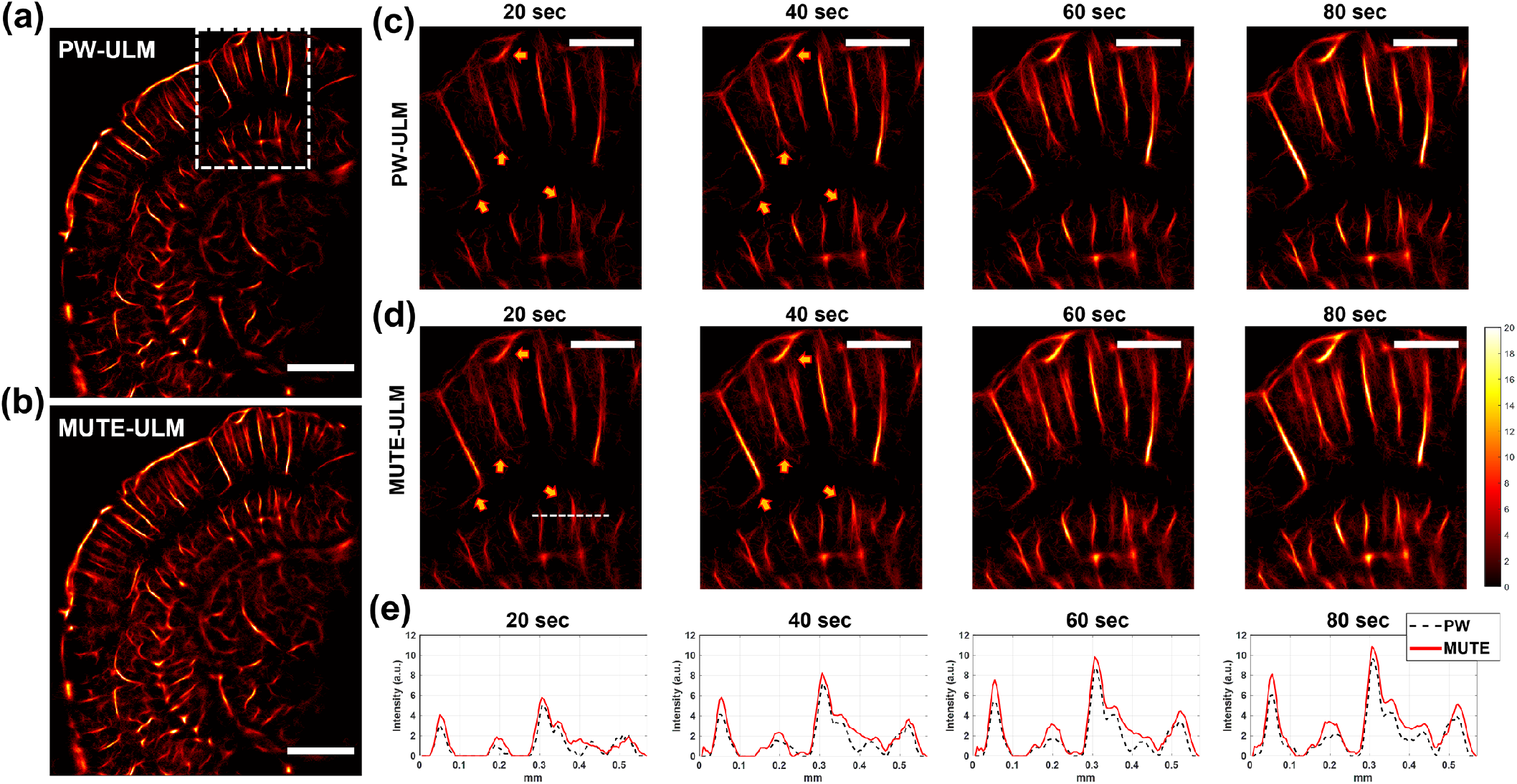
ULM imaging of a mouse brain *in vivo:* (a) PW-, and (b) MUTE-ULM images of mouse brain. Scale bars in (a) and (b) represent 1 mm. Local analysis of (c) PW- and (d) MUTE-ULM images. The images were extracted from the regions indicated by the white box in (a). (e) 1D plots of the vessel signal at different accumulation times from the location indicated by the white dashed line in (d). The yellow arrows indicate different examples of vessels that demonstrate the faster filling speed of MUTE-ULM. Scale bars in (c) and (d) represent 500 μm. Every image was generated in same color scale.

## IV. Discussion

In this paper, we introduced an acoustic analogue to STED for super-resolution ultrasound microvascular imaging. MUTE enhances the performance of ULM by uncoupling overlapping MB signals in space based on acoustic nulls and image subtraction. We demonstrated that MUTE significantly improved the performance of ULM, especially under high MB concentrations. This has strong implications for the practical implementation of ULM because a higher MB concentration leads to a faster vascular filling time, which then translates to shorter data acquisition and higher temporal resolution for ULM.

We first evaluated the performance of MUTE in silico with randomly distributed MBs. We found that MUTE-based localization was highly effective with uncoupling the spatially overlapped MBs that are otherwise inseparable and unlocalizable by conventional ULM. MUTE also contributes to a more accurate MB localization when compared with conventional ULM [Fig. 5].

We then tested the performance of MUTE in vivo on a chicken embryo CAM model and a mouse brain with high concentration MB injections. In both models [Figs. 6 and 7], MUTE-ULM demonstrated significantly faster VF time and higher MB count than conventional ULM. These results suggest that MUTE can effectively mitigate the issue of slow imaging speed of ULM by promoting a more robust MB localization with high efficacy under high MB concentrations, which translates to a faster vascular MB filling rate and an overall reduced data acquisition time.

Unlike in STED where optical fluorophores can be conveniently turned ON or OFF by different wavelength emissions, MBs have a less binary response to acoustic waves and are difficult to transition between the ON and OFF status. Although high MI acoustic pulses can disrupt MBs and switch them OFF, the disruption process takes a long time and even the disrupted MBs (i.e., free gas) can be echogenic. Therefore, in this paper, we resorted to using acoustic nulls and the principles of image subtraction to realize STED for ultrasound super-resolution imaging. Although the concept of using acoustic nulls to enhance ultrasound imaging resolution was reported before (e.g., NSI), these methods exclusively operate in the receive beamforming process and do not actively modulate microbubble response with the acoustic nulls.

For practical implementations of MUTE, there are several tradeoffs that need to be balanced. First, the number of nulls that can be simultaneously generated have a positive relationship with the imaging frame rate: the more the number of nulls that can be simultaneously generated in the FOV to cover a large region, the less the number of transmissions is needed for constructing the full FOV, and hence the higher the imaging frame rate. In this paper, we used 8 elements to generate each null (31 nulls total for the 128-element transducer). The resulting image frame rate was 500 Hz (after 3 angle compounding and 4 lateral translations to cover the full FOV), which is adequate for most ULM applications. The second tradeoff is between the width and the number of the acoustic nulls and their relationship to the imaging frame rate. In theory, a larger number of elements could generate narrower nulls with lower acoustic pressure. However, the more the elements used for each null, the less the number of nulls that can be generated simultaneously and consequently the slower the frame rate. In this study, the width of the acoustic nulls was measured at approximately 50 μm, which corresponds to 1λ. Therefore, MBs with spacing less than 1λ may not get uncoupled effectively. At last, one needs to note that the effect of uncoupling by MUTE is more significant in the lateral direction than in the axial direction. Although steering may introduce uncoupling in the axial direction, due to the limited steering angle that is achievable on conventional ultrasound transducers, MUTE still largely promotes MB separation along the lateral direction. Nevertheless, because the lateral resolution is intrinsically inferior to axial resolution in ultrasound imaging, improvement in the lateral direction is necessary and significant for robust ULM.

## V. Conclusions

In this paper, we demonstrate a new imaging sequence (MUTE – microbubble uncoupling via transmit excitation) that utilizes the principles of acoustic null and image subtraction to uncouple and separate spatially overlapping MB signals that are otherwise unlocalizable in conventional ULM imaging. We showed that MUTE could significantly improve the MB localization efficacy and imaging speed of ULM under high MB concentrations. These results indicate that MUTE has the potential to alleviate the issues of long data acquisition time and low temporal resolution of ULM, which is significant for practical implementations of ULM, especially under clinical settings.

## Notes

### Competing Interest Statement

The authors have declared no competing interest.

